# Deciphering the function of intrinsic and genomics-driven epigenetic heterogeneity in head and neck cancer progression with single-nucleus CUT&RUN

**DOI:** 10.1101/2024.02.14.580230

**Authors:** Howard J. Womersley, Daniel Muliaditan, Ramanuj DasGupta, Lih Feng Cheow

## Abstract

Interrogating regulatory epigenetic alterations during tumor progression at the resolution of single cells has remained an understudied area of research. Here we developed <u>a</u> highly sensitive single-nucleus CUT&RUN (snCUT&RUN) assay to profile histone modifications in isogenic primary, metastatic, and cisplatin-resistant head and neck squamous cell carcinoma (HNSCC) patient-derived tumor cell lines. We find that the epigenome can be involved in diverse modes to contribute towards HNSCC progression. First, we demonstrate that gene expression changes during HNSCC progression can be co-modulated by alterations in both copy number and chromatin activity, driving epigenetic rewiring of cell-states. Furthermore, intratumour epigenetic heterogeneity (ITeH) may predispose sub-clonal populations within the primary tumour to adapt to selective pressures and foster the acquisition of malignant characteristics. In conclusion, snCUT&RUN serves as a valuable addition to the existing toolkit of single-cell epigenomic assays and can be used to dissect the functionality of the epigenome during cancer progression.

## Introduction

Tumour metastasis and acquired drug resistance are key steps in cancer progression that ultimately lead to treatment failure and patient mortality. To identify novel modalities to prevent cancer progression and improve patient survival, it is important to understand the underlying mechanisms that drive these processes. To date, efforts to comprehend the basis of cancer progression have largely focused on uncovering genetic alterations within a tumour population that exert selective fitness, which allows cells to survive during drug treatment. However, cancers can exhibit marked cell-to-cell variation (intra-tumour heterogeneity or ITH) in gene expression and their functional phenotypes that cannot always be explained by mutations or structural variations in their DNA. Non-genetic basis for ITH suggests a potential role for epigenetic mechanisms driving transcriptomic/phenotypic heterogeneity (Brock et al. 2009). Epigenetic alterations offer a heritable mechanism for generating ITH (Easwaran et al. 2014) and occur at greater frequencies in human cancers than genetic mutations (Guo et al. 2019), thereby underscoring the importance of interrogating them when mapping trajectories of cellular plasticity-driven tumour evolution to metastatic or treatment-resistant disease. Cancer cells that survive sub-lethal challenges can often activate stress response pathways which confer early developmental, stem-like features that enable them to cope with further insults (Pisco and Huang 2015). These pro-survival alterations are often manifested in the aberrant modification of histone proteins (Füllgrabe et al. 2011). Furthermore, dysfunction in histone-modifying enzymes have been shown to have a causal relationship with cancer initiation and progression (Wang et al. 2016). Thus, understanding the underlying epigenetic mechanisms that underpin the evolution of cancer is crucial for the discovery of alternative therapeutic interventions to halt or delay its progression and to improve the survival of cancer patients in the clinic.

One such cancer type where epigenetic ITH remains an understudied area of research is head and neck squamous cell carcinoma (HNSCC). Single cell studies on HNSCC mainly focused on transcriptomic ITH, underscoring the need to investigate epigenetic control of ITH in HNSCC (Puram et al. 2017; Quah et al. 2023; Choi et al. 2023; Qi et al. 2021). An area where epigenetic ITH can have a role in HNSCC progression is in the alterations of histone modifications. Histone modifications can occur rapidly as cells respond and adapt to the environment. Classically these changes are identified by chromatin immunoprecipitation followed by sequencing (ChIP-seq). However, classical ChIP-seq is only suitable for bulk cell assays, which have limited application for discerning epigenetically distinct subpopulations within tumours. This makes it difficult to determine whether plasticity or lineage transitions are a result of Darwinian selection of rare, pre-existing clones; or adaptation, where dynamic epigenetic changes may activate distinct transcriptomic programs that lead to the emergence of new phenotypes. Thus, there is a need for studies employing methodologies that can detect histone modifications at the resolution of single cells in order to answer questions related to how epigenetic heterogeneity and plasticity may drive cell-state transitions during HNSCC progression. Importantly, given that many readers and writers of different histone marks are amenable to pharmacological interventions (Helin and Dhanak 2013; Kakiuchi et al. 2021), a greater understanding of their regulatory function during tumour <u>progression</u> could result in the identification of new therapeutic intervention strategies for HNSCC patients.

Methods for profiling histone modifications or transcription factors in single cells generally involve either immunoprecipitation, using an antibody immobilized nuclease or an antibody immobilized transposase – as exemplified by scChIP-seq, uliCUT&RUN and scCUT&Tag, respectively (Grosselin et al. 2019; Hainer et al. 2019; Kaya-Okur et al. 2019). Due to the paucity of DNA within single cells and the implicit requirement for selection of only a tiny fraction of this material, these methods and others tend to have low numbers of filtered reads and/or poor specificity. Single cell ChIP-seq methods employ microfluidic devices to compartmentalize single cells, and although thousands of cells can be analyzed simultaneously the number of unique reads per cell tends to be low due to inefficient reactions within individual droplets. Additionally, the requirement for special equipment limits the adoption of this approach by the wider scientific community. On the other hand, the use of immobilized transposases has generated significantly more traction (Ai et al. 2019; Carter et al. 2019; Wang et al. 2019), largely because these methods do not require a library preparation stage that normally leads to additional loss of already scarce material. However considerable off-target transposase binding occurs without stringent salt washes, which can inadvertently detach proteins of interest from DNA unless they are tightly bound (Kaya-Okur et al. 2020). CUT&RUN was developed as an alternative to ChIP-seq that exhibits significantly less background noise (Skene and Henikoff 2017), which enables it to profile as few as a 100 cells (Skene et al. 2018). A single cell version - uliCUT&RUN - was demonstrated to localize NANOG and SOX2 transcription factors (TF) in rare populations of mouse embryonic stem cells (Hainer et al. 2019). However the extensive sequencing depth required for this method makes it impractical for analyzing large numbers of single cells (Patty and Hainer 2021).

In this manuscript, we developed single nucleus CUT&RUN (snCUT&RUN) assay to profile histone modifications in single nuclei of HNSCC tumour cells. Interrogating isogenic patient-derived cell lines (PDC) representing primary, metastatic and cisplatin-resistant tumour cell lines from HNSCC (Chia et al. 2017), we found that H3K4me3 expression remains relatively stable across the distinct evolutionary states. In contrast, H3K27ac modifications were more pronounced and exhibited more divergence corresponding with global changes in acetylation in progressed tumour states. Overall, the same cell types from different patients displayed unique epigenetic profiles – indicating distinct events may have led to the development of the primary tumour, and subsequently their progression to metastatic and drug resistant states. Notably, snCUT&RUN inferred the presence of copy number variations in regions with histone modifications. We found that specific differences in histone modification can be accentuated by genetic copy number alterations (CNA) between cancer cells, suggesting that modulation of the cellular epigenome by genetic aberrations could represent an additional mechanism for cancer progression. Furthermore, we discovered that intratumour epigenetic heterogeneity (ITeH) may give rise to subpopulations within primary cells that mimic the metastatic and/or drug-resistant cell states. This subset of cells, undetected by bulk approaches, may be epigenetically primed to transition to a progressed state. H3K27ac modification in single cells of the progressed states displayed stress response signatures, suggesting that dynamic alterations of the epigenome upon drug insults could underlie adaptational changes in their transcriptome and phenotype. Altogether, we demonstrate that high resolution profiling of histone modifications in single cells with snCUT&RUN can yield valuable insights into epigenetic heterogeneity in HNSCC, complementing existing single cell transcriptomic studies.

## Results

### Development and performance of single nucleus CUT&RUN

Tagmentation-based methods for single cell histone modification profiling (e.g., scCUT&TAG) have limitations with efficiency for the following reasons: 1) two independent transposition reactions are needed to create a sequenceable fragment, 2) productive reactions require that the adapters introduced by transposition on each end of the fragment to be of different “types” – which only happens with 50% probability. As such, the median unique read counts from methods such as scCUT&Tag tend not to exceed 10,000 which limits the detection of subtle epigenetic changes in many applications. Considering these challenges, we focused on the nuclease-based protocol – CUT&RUN. Unlike transposon-based methods which get expended during the reaction, a single pA-MN enzyme in CUT&RUN can catalyze fragmentation of DNA on both sides of a given nucleosome. This means that after digestion, all fragments have the potential to yield sequenceable product after the library preparation stage. Although the library preparation protocol is more complex compared to scCUT&TAG, we believe that the increased sensitivity of snCUT&RUN would better cater to the needs for applications that require higher-sensitivity histone profiling at single cell resolution.

A schematic of snCUT&RUN is shown in Figure 1A, and the detailed protocol is provided in the Materials and Methods section. The original CUT&RUN method has low background noise and requires a fraction of the sequencing depth compared to ChIP-seq. We found that several enhancements could transform this technique into a single cell method that could be performed in a standard molecular biology laboratory with access to a FACS facility (or other means of single cell isolation). First, we found that it was imperative to maintain nuclear integrity throughout the entire workflow to minimize background reads and ensure high quality data. Damage to nuclei results in release of ambient DNA, clumping of nuclei and loss of material. Hence, we formulated a lysis buffer with minimal detergent concentration; and the addition of sucrose plus BSA resulted in lysis with vastly reduced clumping when resuspending pellets during washing steps throughout the workflow. Lysis, antibody, and pA-MN binding steps were all performed in bulk, which reduced the amount of handling and the loss of individual cells after isolation. Second, directly after pA-MN incubation, the nuclei were stained in buffer containing a nuclear dye plus a labelled anti-nuclear pore complex antibody. This step greatly improved the efficiency in isolating single intact nuclei rather than multiple nuclei or debris. Third, low salt conditions have been shown to prevent pA-MN from diffusing after digestion, thereby reducing off-target cleavage of background DNA (Meers et al. 2019). We adopted this approach by directly sorting single nuclei into a low-salt calcium buffer to initiate nuclease activity. Lastly, given that the library preparation stage is the major source of sample loss, inactivation of pA-MN, end-repair, A-tailing, indexed adapter ligation and SPRI steps were all performed with no tube transfers. The amount of paramagnetic SPRI beads used to remove unbound adapters were doubled relative to PEG/NaCl to increase the surface area available for binding to the DNA template. Furthermore, 50N neodymium magnets were used for separation to minimize dead volumes. By avoiding proteinase K treatment, intact nuclei and high molecular weight debris could be observed being retained by the SPRI beads after eluting the target DNA, thereby reducing background. PCR was performed on pooled samples, the products concentrated via precipitation, and adapter monomers and dimers removed with a second set of SPRI bead treatment. The template from 192 libraries were then sequenced on a single MiSeq chip (Methods).

**Fig. 1.**
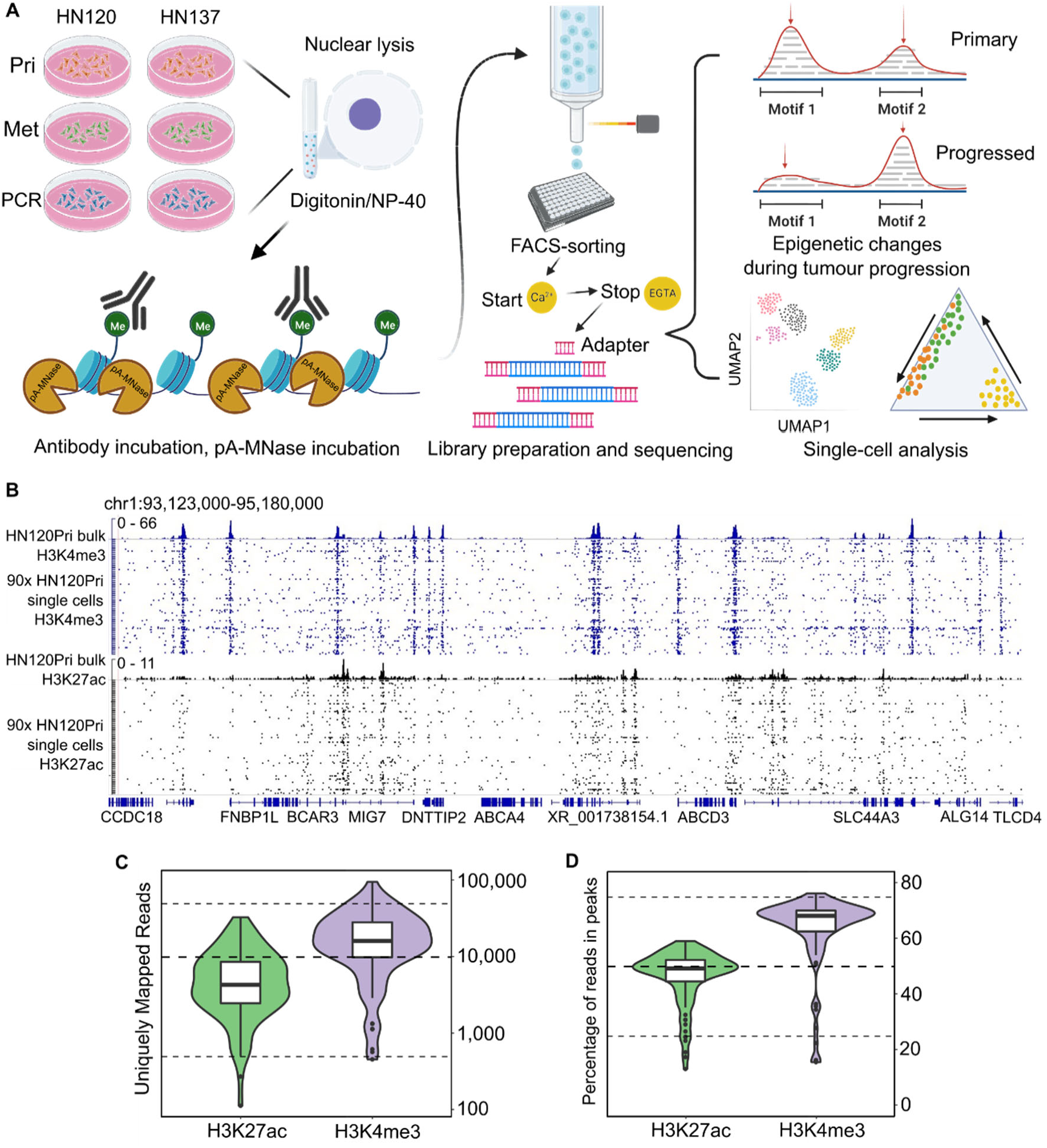
Schematic representation of overall workflow and quality control of snCUT&RUN. **(A)** Schematic overview of snCUT&RUN, applied on matched primary, metastatic, and primary cisplatin-resistant patient derived head and neck cancer cell lines from two patients (HN120 and HN137). **(B)** Representative IGV-track image showing H3K4me3 and H3K27ac single-cell profiles of 90 HN120Pri cells, with corresponding bulk cell data for each mark. **(C)** Violin- and boxplots showing the distribution of the number of unique mapped reads (UMRs) for each single-cell, for both H3K4me3 (median = 16,296) and H3K27ac (median = 4352). Dotted lines are at 50,000, 10,000 and 500 UMRs. **(D)** Violin- and boxplots illustrating the percentage of reads in peaks for each single cell. H3K4me3 median = 68%, H3K27ac median = 49%. Dotted lines are at 75, 50 and 25 percent.

We benchmarked the technical performance of snCUT&RUN on 90 sorted nuclei, probed either with anti-H3K4me3 or anti-H3K27ac antibodies. The sequencing read distribution of single cells exhibited a high degree of similarity with bulk cell populations probed with the respective histone-specific antibodies (Fig. 1B), demonstrating excellent specificity of our protocol. H3K4me3 is a histone mark that is generally associated with active promoters. The read density profile from H3K4me3 snCUT&RUN samples showed similar profiles around transcriptional start sites (TSS) to that of the bulk cell set; with corresponding patterns of nucleosome depletion (Supplemental Fig. S1A), indicating the high resolution of this assay. Most strikingly, the median number of unique reads for the H3K4me3 was 16,296, whereas the corresponding number for H3K27ac was 4,352. As an indication of signal-to-noise, the median Fraction of Reads in Peaks (FRiP) were 68% for H3K4me3 and 49% for H3K27ac (Fig. 1D), which to the best of our knowledge are on par with other single cell histone assays.

Comparison of the histone profiles of pooled snCUT&RUN with bulk CUT&RUN yielded a Pearson correlation 0.956 and 0.6089 for H3K4me3 and H3K27ac, respectively (Supplemental Fig. S1B and 1C). Collectively, these data indicated that snCUT&RUN recapitulated bulk CUT&RUN data to a high degree and that this method has the potential to resolve subtle epigenetic differences that might be expected within cancer cell populations. Once over 1,000 cells had been screened for H3K4me3 and H3K27ac, we compared the performance of snCUT&RUN with previously published data on scCUT&Tag (Kaya-Okur et al. 2019) by ranking the cells by number of UMRs (Supplemental Fig. S1D). Although different antibodies were used in the two methods, we reasoned that there should be considerable overlap between the profiles of H3K4me2 and H3K4me3 modifications; being present at transcribing, and poised plus transcribing genes, respectively (Hyun et al. 2017). Under our experimental conditions, we could achieve almost an order of magnitude more uniquely mapped reads than scCUT&Tag. As a second metric we compared snCUT&RUN with previously reported single cell assay using CUT&RUN (uliCUT&RUN) by determining the percentage of reads which map uniquely (Supplemental Fig. S1E). Notably, snCUT&RUN produced better mapping rates after deduplication (12% for H3K27ac and 13% for H3K4me3) compared to 0.3-0.7% with uliCUT&RUN (Patty and Hainer 2021). This could be attributed to loss of material with uliCUT&RUN, where low template concentrations during PCR can cause an increase in artifacts (Ruiz-Villalba et al. 2017).

### Copy number amplifications may drive epigenetic reprogramming during metastatic progression in HNSCC

Having established the technical validity of snCUT&RUN, we utilized the R packages Signac and Seurat (Stuart et al. 2019, 2021) for an integrated analysis of snCUT&RUN data on previously established isogenic patient-derived HNSCC models that represent primary (Pri), metastatic (Met) and cisplatin-resistant primary tumor (PCR) cell lines from two individual patients (HN137 and HN120 (Chia et al. 2017; Sharma et al. 2018)) (Supp. Table 1). We used a set of filtering parameters (described in detail in Methods) to eliminate low quality cells. After filtering, single-cell profiles of 1,107 and 1,048 cells for H3K4me3 and H3K27ac-specific ChIP data respectively were used for downstream analysis. UMAP embedding of H3K4me3 and H3K27ac showed slightly different patterns and varying degrees of overlap between the cell types (Fig. 2A). From H3K4me3 profiles, HN120Pri and HN120Met cells were indistinguishable, and a similar observation was made when comparing HN137Pri and HN137PCR cells (Fig. 2A, left).

**Fig. 2.**
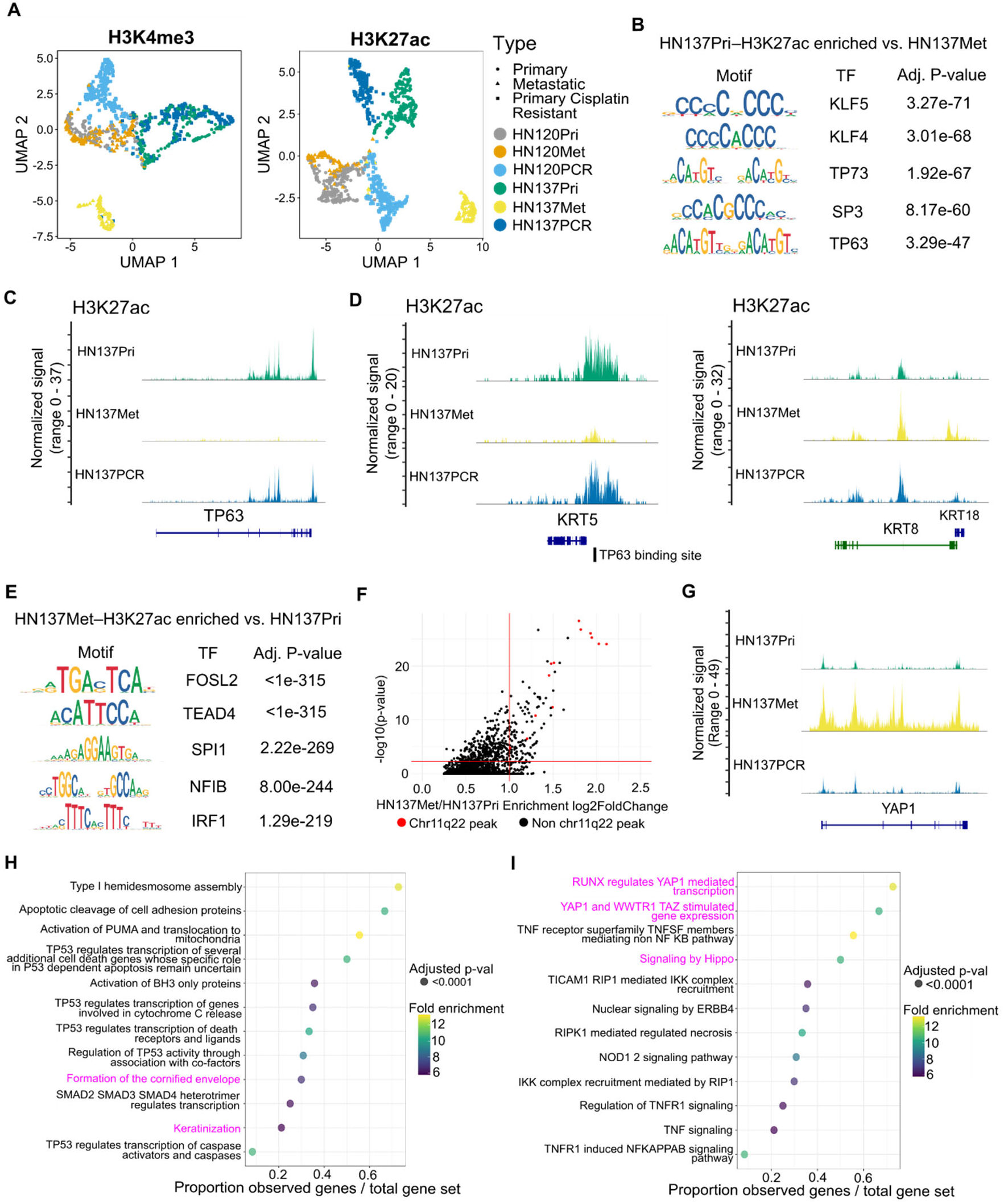
Epigenomic changes during HNSCC progression suggest distinct, patient-specific epigenetic drivers of tumour evolution. **(A)** UMAP embedding for H3K4me3 (left) and H3K27ac (right) for HN120 and HN137 single-cells. **(B)** Enriched transcription factor motifs for HN137Pri derived from H3K27ac peaks. **(C)** Coverage plot of H3K27ac signal at the TP63 locus, showing loss of H3K27ac in HN137Met. **(D)** Coverage plots of H3K27ac signal at the KRT5 (left) and KRT8/18 (right) loci. TP63 binding site near the KRT5 promoter is indicated. **(E)** Enriched transcription factor motifs for HN137Met. **(F)** Top enriched H3K27ac peaks in HN137Met, with peaks at the chr11q22 locus containing YAP1 highlighted in red. **(G)** Coverage plot of H3K27ac signal at the YAP1 locus. **(H, I)** Dot plot indicating the Reactome pathways enriched in HN137Pri **(H)** and HN137Met **(I)**. Highlighted in magenta are terms indicating YAP1-mediated loss of differentiation in HN137Met.

In contrast, the PDCs were organized into more distinct clusters when H3K27ac profiles were used for clustering (Fig. 2A, right), suggesting that H3K27ac expression (active promoters and enhancers) is more divergent between primary and progressed cell states compared to promoter activity alone. Notably, HN120PCR and HN137Met seemed to have distinct profiles for both H3K4me3 and H3K27ac, indicating significant differences in their chromatin states when compared to other PDCs. We confirmed our clustering results by calculating Pearson-correlations between the H3K27ac/H3K4me3 profiles of the different cell types (Supplemental Fig. S2A,B). Furthermore, we did not detect significant batch effects in our clustering (Supplemental Fig. S2C,D).

Next, we sought to infer how changes in epigenetic landscape could drive HNSCC progression using the snCUT&RUN data. After Signac-integrated peak calling, we obtained a set of 606,594 H3K27ac and 391,115 H3K4me3 peaks present in at least one cell line (Supplemental Fig. S3A,B). Out of these peaks, most are either cell-line specific or shared between all cell lines. However, snCUT&RUN was able to capture peaks that were shared by primary tumour and matched progressed cell lines (Supplemental Fig. S3C,D). Amongst the primary tumour/progressed paired cell lines with the starkest H3K27ac/H3K4me3 profile differences are HN137Pri and HN137Met (Fig. 2A). Transcription factor (TF) motif binding analysis in HN137Pri revealed an enrichment of TP63, a key lineage-determining regulator of epidermal keratinocyte identity (Qu et al. 2018), and KLF4, which was previously shown to regulate differentiation in the basal layer of the oral epithelia (Segre et al. 1999) (Fig. 2B). Upon plotting the H3K27ac coverage signal, we observed near loss of H3K27ac at the TP63 locus indicating downregulation of TP63, and thereby loss of lineage fidelity, in HN137Met (Fig. 2C). Since TP63 is known as a key regulator of basal keratinocyte identity, we also focused on the keratin (KRT) loci and found a near loss and gain of H3K27ac at the KRT5 and KRT8/18 loci, respectively, in HN137Met (Fig. 2D). KRT5/KRT18 immunofluorescence (IF) corroborated our findings from snCUT&RUN data (Supplemental Fig. S4A). Furthermore, we confirmed the downregulation of TP63 and KRT5 and upregulation of TEAD4 and KRT8/18 during HNSCC metastasis, with the analysis of two previously published scRNA-seq datasets from primary and metastatic HNSCC (Puram et al. 2017; Sharma et al. 2018) (Supplemental Fig. S4B-J). We found an annotated TP63 binding site near the KRT5 promoter, supporting the notion of KRT5 expression being driven by TP63. Additionally, H3K27ac profiles of the KRT5 and KRT8/18 loci in HN120 PDCs corroborated the immunofluorescence-based protein expression (Supplemental Fig. S5A). Moreover, we observed concordant H3K4me3 activity at the KRT5/8/18 loci of HN120 and HN137 PDCs (Supplemental Fig. S5B). Hence, the simultaneous presence (or absence) of H3K4me3 and H3K27ac is a strong indicator of epithelial cell identity. The findings of activating epigenetic marks in the KRT8/18 locus in HN137Met is significant, since high KRT8 expression has been associated with detachment of cells from tumours and seeding of lymph node metastasis in HNSCC (Matthias et al. 2008).

The TFs enriched in HN137Met include members of the TEAD family such as TEAD4, a downstream activator in the Hippo pathway, and FOSL2, a regulator in cellular differentiation (Fig. 2E). Visualization of transcription factor activities at single cell level showed that loss of TP63 and TEAD4 in HN137Met and HN137Pri respectively (Supplemental Fig. S5C). Previously, it was shown that TEAD can repress TP63 promoter activity and protein expression (Valencia-Sama et al. 2015). We further assessed the differential peaks between HN137Met and HN137Pri. Strikingly, the top enriched peaks in HN137Met were predominantly located at the chr11q.22 locus containing YAP1 (Supplemental Fig. S3F). Increased activity of YAP1, in conjunction with its binding partner TEAD, was previously shown to promote proliferation and metastasis in multiple cancers, including HNSCC (Lamar et al. 2012; Chia et al. 2017; Omori et al. 2020). We also found a significant increase in H3K27ac activity of YAP1 (Fig. 3G). YAP1 is frequently amplified in HNSCC and indeed, we have previously reported YAP1 to be amplified in HN137Met, compared to the patient-matched HN137Pri (Chia et al. 2017; Shin and Kim 2020).

**Fig. 3.**
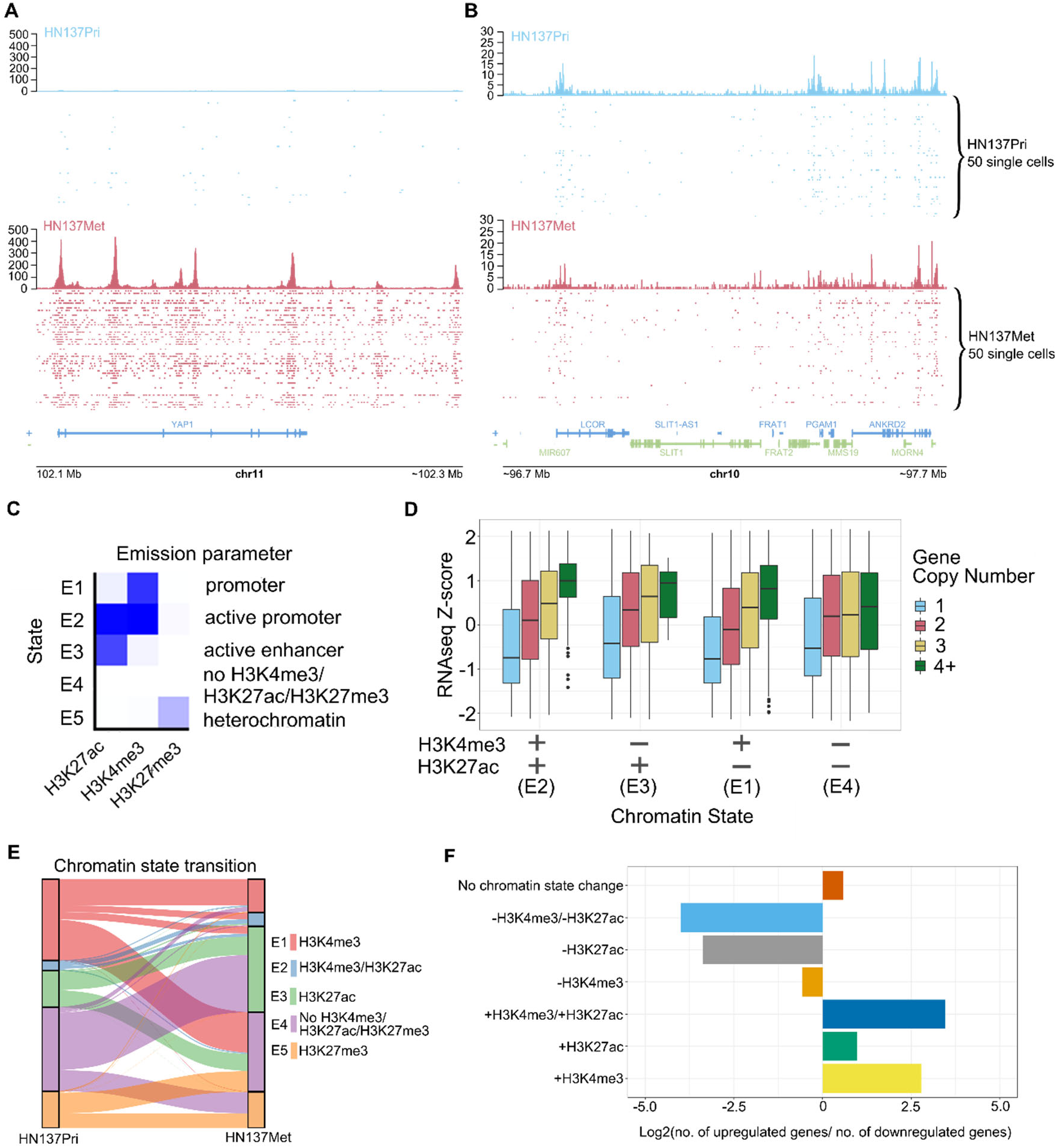
Regulation of gene expression through the interaction between copy number variations and chromatin state. **(A)** Distribution of unique H3K27ac reads in single cells at the YAP1 locus in HN137Pri (blue) and HN137Met (red). Bulk H3K27ac signal and H3K27ac signal in single cells are shown. **(B)** Distribution of unique H3K27ac reads in a random, non-copy number different locus. **(C)** ChromHMM results identifying 5 chromatins states consisting of combinations of H3K4me3, H3K27ac, and H3K27me3 modifications in HN120 and HN137 PDCs. **(D)** Boxplots representing RNA-seq Z-scores of HN137Met genes, stratified by both gene CN as well as chromatin state annotation. **(E)** Alluvial plot showing chromatin state transition between HN137Pri and HN137Met. Unmodified chromatin (E4, purple) which remained unmodified during metastatic progression was excluded from the plot for visibility purposes. **(F)** Log2 ratio of the number of upregulated (Log2 FC > 1, p < 0.001) genes to the number of downregulated (Log2 FC < −0.5, p < 0.001) genes per chromatin state change during metastatic progression of patient HN137 (HN137Pri > HN137Met).

Finally, we assessed top differential enriched peaks (p < 0.005) in HN137Pri and HN137Met and used the R package rGREAT (McLean et al. 2010; Gu 2022) to look for enriched biological pathways. Top enriched Reactome pathways in HN137Pri related to TP53 activity (a member of the same family as TP63) and keratinocyte differentiation, while top pathways in HN137Met include activation of the Hippo pathway, which includes YAP1 and TEAD, and TNF-mediated activation of NFκB (Fig. 2H,I). Altogether, these results suggest that H3K4me3/H3K27ac profiles derived from snCUT&RUN could be used to confirm phenotypical changes occurring between primary and progressed HNSCC. Furthermore, the results indicate that considerable changes in epigenetic modifications could be associated with copy number amplifications, and that a singular focal amplification, such as what was observed with YAP1, could lead to epigenetic reprogramming of HNSCC cells to acquire metastatic capabilities.

### Genetic-epigenetic alterations correlate with changes in gene expression

Since our data suggested that genetic drivers such as YAP1 amplification could lead to changes in the epigenome, we further explored how changes in gene copy number and the combination of active histone marks, such as H3K4me3/H3K27ac, could correlate with alterations in gene expression. We first visualized H3K27ac signal of single cells at the YAP1 locus for HN137Pri and HN137Met and confirmed our previous findings from Signac in which we observed a marked increase in H3K27ac activity in the YAP1 signal in HN137Met (Fig. 3A). In contrast, regions without copy number differences do not exhibit significant differences in H3K27ac signal (Fig. 3B). These observations reflect the copy number amplification at the YAP1 locus in HN137Met. Thus, snCUT&RUN can reveal not only the epigenetic changes, but also points towards regions potentially affected by copy number alterations. Several other loci with potential differences in copy number were also detected (Supplemental Fig. S6).

Next, to correlate chromatin profiles with gene expression and copy number variations, we performed bulk RNAseq and whole exome sequencing (WES) on all PDCs. DNA copy number variation has been reported to correlate with gene expression changes. Indeed, we observed a clear positive correlation between PDC gene expression Z-score and the copy numbers for the same genes (Supplemental Fig. S7A). These results corroborate the notion that changes of gene expression in key cancer genes may arise from copy number variations (CNV). Nevertheless, we hypothesized that epigenetic modifications in such genetically altered loci may further modulate the expression of genes that reside within the same loci. This effect could be compensatory (epigenetic silencing of an amplified gene or epigenetic activation of a deleted gene) or conversely it could accentuate the effect of CNV (epigenetic activation of the amplified gene or silencing of the remaining copy of a deleted gene). To investigate these hypotheses, we first used ChromHMM (Ernst and Kellis 2012) to perform joint analysis based on pseudobulk H3K4me3, pseudobulk H3K27ac from snCUT&RUN as well as bulk H3K27me3 CUT&RUN data. ChromHMM results revealed five chromatin states, reflecting distinct chromatin state annotations: weak promoter (E1, H3K4me3 only), active promoter (E2, H3K4me3+/H3K27ac+), active enhancer (E3, H3K27ac only), heterochromatin (E5, H3K27me3 only) and unmodified regions with none of the probed chromatin marks present (E4) (Fig. 3C). We then combined gene copy number (CN), chromatin state annotation, and gene expression data to investigate how chromatin state and gene CN may synergistically influence gene expression. The results showed that in the presence of activating epigenetic marks, gene expression Z-scores increase with copy number, and gene expression levels are higher (Fig. 3D, Supplemental Fig. S7B). Genes that have simultaneous H3K4me3 and H3K27ac marks meanwhile have higher gene expression Z-scores compared to singly-marked genes. Meanwhile, genes that do not have an activating epigenetic mark surprisingly show little to no correlation with gene expression Z-score despite their CN status. Our results therefore underscore the importance of epigenetic modifications to modulate the effects of CNVs on gene expression during HNSCC progression. Without the appropriate epigenetic modifications, the effects of CNV are masked and would not manifest in gene expression. These results further highlight the need to consider both CNVs as well as epigenetic modifications on gene regulatory elements to understand factors influencing the progression of HNSCC more comprehensively and accurately.

Next, we decided to interrogate how epigenetic profiles at single cell resolution could assess the function of chromatin state dynamics in driving HNSCC progression from primary cancers to a progressed state (metastatic or treatment resistant). We therefore analyzed genome-wide chromatin state transitions between paired, patient-matched primary and progressed PDCs (HN120Pri to HN120Met, HN120Pri to HN120PCR (drug resistant), HN137Pri to HN137Met, and HN137Pri to HN137PCR). We found that most of the epigenome (∼95%) remained devoid of H3K27ac, H3K4me3 and H3K27me3 (E4). Furthermore, we observed diversity in histone mark changes across the various transitions (Fig. 3E, Supplemental Fig. S8A). The general trend, however, was that comparing primary tumour PDCs with their matched progressed PDCs, there appeared to be a greater proportion of genomic regions that acquire the active H3K27ac mark in the progressed state. Meanwhile, the proportion of H3K4me3-marked regions either decreased or remained unchanged. Previous reports have suggested that alterations in cellular metabolism during cancer progression can modulate the levels of co-factors that are substrates of chromatin-modifying enzymes (e.g., acetyl-CoA for histone acetyltransferase) consistent with our observation of increased H3K27ac marks in progressed state. Metabolic reprogramming in cancer cells have been shown to promote EMT by epigenetic activation of EMT-associated genes (Wang et al. 2020). Indeed we found increased H3K27ac signal at the SERPINE1 and SERPINA1 genes, which were previously associated with metastasis, apoptosis resistance, and poorer prognosis in HNSCC (Supplemental Fig. S8B) (Pavón et al. 2015), supporting the epigenetic basis of metastasis in HNSCC.

Finally, we investigated whether significant changes in gene expression during HNSCC progression correlated with changes in chromatin state. We filtered the top downregulated (log2 fold change < −0.5, p < 0.001) and upregulated genes (log2 fold change > 1, p < 0.001) during the HN137Pri > HN137Met transition and calculated the enrichment of gene expression change per chromatin state transition. We found that downregulation of gene expression was indeed associated with the loss of H3K4me3/H3K27ac active marks, whereas genes displaying upregulated expression were associated with a gain of either or both H3K4me3/H3K27ac marks (Fig. 3F). Altogether, the data suggests that during HNSCC progression, the gain of H3K27ac in key enhancers as a consequence of a global increase in acetylation may serve as the primary drivers of changes in gene expression that promote cell-state transitions. Furthermore, our data also suggests that alterations in chromatin states occurring between primary and progressed HNSCC are dynamic and heterogeneous, reflecting the diversity of epigenetic changes during tumour progression. These results further corroborated our previous findings that global changes in the epigenome could lead to change in gene expression, highlighting the need for considering chromatin state changes when exploring mechanisms of HNSCC progression. Finally, these results collectively suggest that changes in gene CN and chromatin states may be interconnected in HNSCC, and that multimodal integration of genomic and epigenomic data would be essential to generate a more comprehensive understanding of HNSCC progression.

### Epigenome profiling at single nuclear resolution identifies subpopulations of primary HNSCC tumour cells with higher propensity for metastatic progression

Our results above reveal that both genetic changes, exemplified by CNV, and epigenetic flux work in tandem to rewire the gene expression network that drives cancer progression. Amplifications on key oncogenes (e.g., YAP1) that associate with crucial transcription factors could have a disproportionate effect on changing the global epigenetic landscape through interactions with histone writers. We speculate that such structural alterations could account for why the epigenetic profiles of HN137Met or HN120PCR were found to be far removed from their primary counterparts. On the other hand, the transitions between HN137Pri → HN137PCR and HN120Pri → HN120Met did not reveal clear genetic drivers of progression identified through whole exome sequencing analysis. In the context of HNSCC progression, some cell subpopulations, especially those at the invasive borders of primary tumors, have been shown to express a partial epithelial-to-mesenchymal (pEMT) transcription program (Puram et al. 2017). Compared to irreversible genetic changes, non-genetic changes could be much more plastic and heterogeneous among cancer cells. Microenvironmental pressures may lead to primary tumour cancer cells to adapt by acquiring a range of epigenetic states and this cellular plasticity could confer survival and growth advantage under the selective pressure of metastasis or drug treatment. We therefore sought to utilize the snCUT&RUN data to explore ITeH as a factor driving HNSCC progression in absence of a clear genetic driver. We first analyzed the transition between HN137Pri and HN137Met, since these PDCs were found to have higher similarity in H3K4me3/H3K27ac profiles as well as phenotype (Fig. Supplemental Fig. S2A, Supplemental Fig. S4A).

Analyzing H3K27ac profiles, we found a lower number of differentially enriched peaks (p-value < 0.005) in HN137PCR versus HN137Pri (2,245), compared to HN137Met versus HN137Pri (4,797). Hence, changes in H3K27ac expression during the cisplatin resistance of HN137Pri appeared to be less widespread compared to the metastatic progression of patient HN137. These results point to a gradual change in the epigenome caused by ITeH, rather than a genetically driven change during progression to a drug-resistant state. To further investigate ITeH in the cisplatin resistance progression of HN137Pri, we defined PDC-specific “modules”, which are features consisting of the top fifty differential peaks defining a specific PDC-state. This analysis was inspired by scRNA-seq analysis, where modules are defined as groups of genes that are part of the same module/program - typically referring to cell-type specific gene expression signatures. After defining H3K27ac modules for each PDC, we then computed ChromVAR deviation Z-scores of each module for each individual cell. In our case, the Z-score represents how far the H3K27ac profile of that cell deviates from the average H3K27ac profile of every cell within the peaks as defined by the PDC-specific modules. Analyzing the ChromVAR deviation Z-scores of the HN137 isogenic cell lines, we found unsurprisingly that each PDC has the highest module score of their individual module (e.g., HN137Met has highest Z-score for the HN137Met module) (Fig. 4A). The deviation Z-score profiles confirmed that each module is capable of distinguishing identities of individual PDCs. However, we also observed a degree of variability in Z-scores within a given cell type, prompting the question whether this represents the possibility of epigenetically heterogeneous state within the PDCs. We therefore normalized the primary tumour, metastatic, and primary tumour cisplatin resistant module scores of each single cell and plotted the normalized scores using ternary plots, making use of the three-variable nature of our data to visualize the ratio of Pri/Met/PCR module score per single cell (Fig. 4B). Ternary plot visualization confirmed our previous findings, showing that HN137Pri and HN137PCR cells are in a continual axis based on their module scores. Such continuity was not observed between HN137Pri and HN137Met, suggesting that the HN137Pri → HN137PCR transition is characterized by intrinsic epigenetic intratumoural heterogeneity and not defined by strong genetic/epigenetic drivers, as is the case with HN137Pri → HN137Met progression (Fig. 4B).

**Fig. 4.**
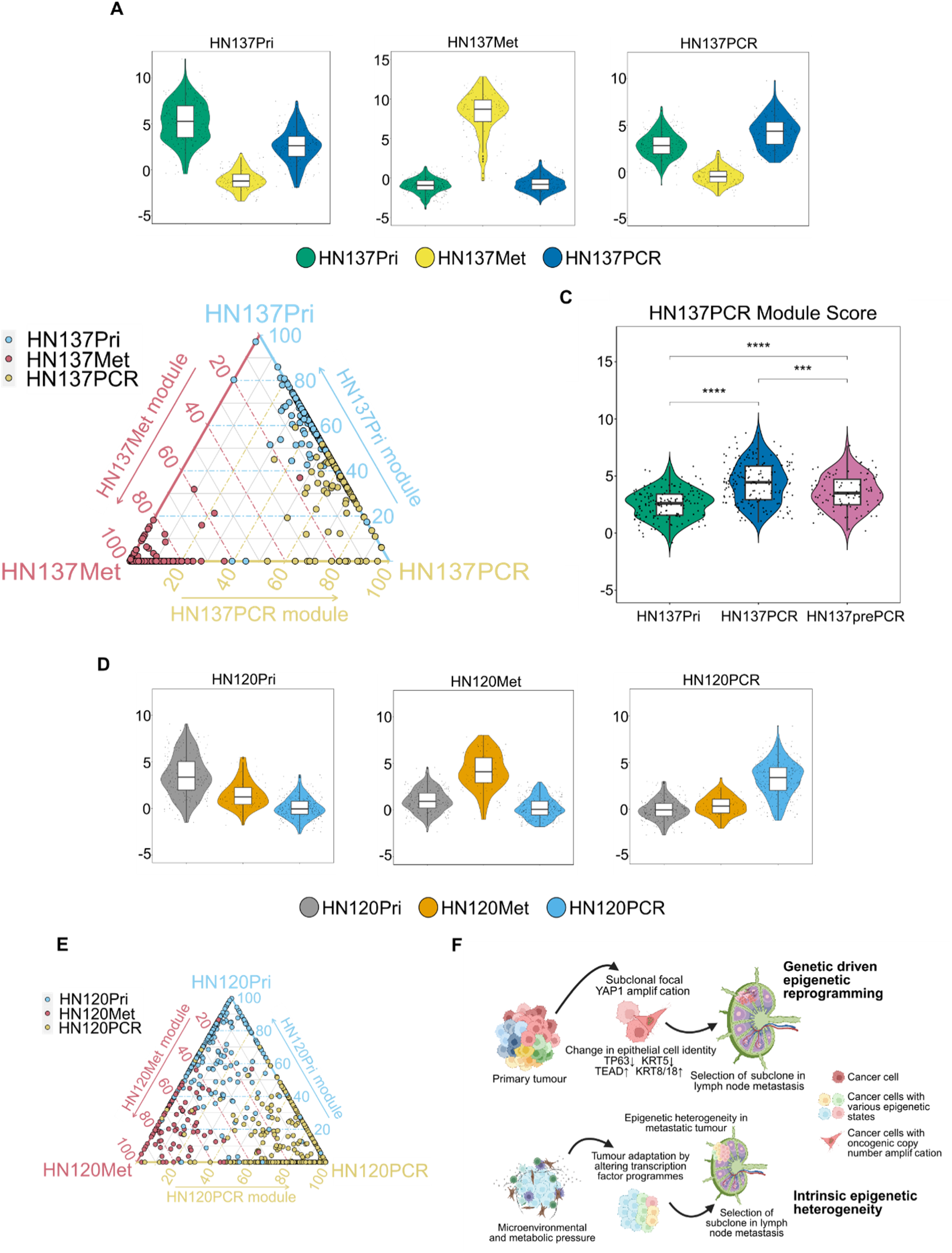
Epigenetic heterogeneity-driven HNSCC progression. **(A)** PDC-specific HN137 H3K27ac module score. Each module consists of the top 50 peaks for a particular PDC, and each module score was back calculated for each single-cell using Signac’s AddChromatinModule function. **(B)** Ternary plot showing the ratios of Pri/Met/PCR module scores for patient HN137 **(C)** Violin plot depicting the recalculated HN137PCR module score after identification of the HN137prePCR subpopulation (*** p-value <= 0.001, **** p-value <= 0.0001). **(D)** PDC-specific HN120 H3K27ac module score. **(E)** Ternary plot showing the ratios of Pri/Met/PCR module scores for patient HN120. **(F)** Model of diverse ways in which epigenetic changes can lead to adaptation of cellular phenotype to acquire a more aggressive state.

To further investigate epigenetic heterogeneity, we set apart HN137Pri cells with Pri/Met/PCR module score ratio of >0.4 Pri and >0.4 PCR and annotated these cells “HN137prePCR” cells. We obtained 93 HN137Pri cells (representing 37% of the primary population) fulfilling this criterion. HN137prePCR cells were found to have a higher average score of the HN137PCR module compared to the remaining HN137Pri cells, yet still below the average of HN137PCR cells (Fig. 4C). This could potentially indicate that HN137prePCR cells represent a primary tumour cell subpopulation which is predisposed towards becoming cisplatin resistant because of their “cisplatin resistant-primed” epigenetic state. To exclude UMR count and FRiP as confounding factors of the module scores, we compared the UMR count and FRiP of HN137prePCR cells against the remaining HN137Pri cells, as well as HN137PCR cells. HN137prePCR cells did not have higher UMR count on average as compared to the remaining HN137Pri as well as HN137PCR cells (Supplemental Fig. S9A). Furthermore, HN137prePCR cells did not have higher FRiP values as compared to HN137PCR and HN137Pri cells (Supplemental Fig. S9B). Hence, UMR count and FRiP were not deemed to be significant factors in determining module scores.

Finally, we investigated whether ITeH could drive the lymph node metastatic progression of patient HN120 (Fig. 4D). In a similar approach to investigate whether ITeH drives the progression of cisplatin resistance in patient HN137, we isolated HN120Pri cells with >0.4 HN120Pri H3K27ac module ratio and >0.4 HN120Met module ratio. We observed 26 cells that met this criterion (∼9% of the total HN120Pri population) and relabeled these HN120Pri cells as the “HN120preMET” subpopulation (Fig. 4D, E). Similar to HN137prePCR, recalculation of the HN120Met module score showed that the HN120preMET cells have higher average deviation Z-score for the HN120Met module as compared to the remaining HN120Pri cells, but still lower than HN120Met cells (Supplementary Fig. 9C). Overall, the HN120preMET and HN137prePCR analyses support the existence of H3K27ac landscape heterogeneity in HNSCC tumours and suggests that in the absence of strong genetic drivers, tumour adaptation could drive ITeH in primary HNSCC tumours, leading to certain selected, epigenetically primed, subpopulations acquiring a phenotype with higher propensity to progress into a more malignant state.

## Discussion

Intratumour heterogeneity of chromatin states can serve as an epigenetic driver of HNSCC progression. However, studies using single-cell methods to profile epigenetic heterogeneity in HNSCC models of cancer progression remain lacking. Here we developed snCUT&RUN, a robust method to profile histone modifications at the single-cell level. We showed that snCUT&RUN could provide insights in chromatin state transitions between isogenic primary and progressed HNSCC PDCs at the level of both global and single-cell resolution. At the global level, we found that generally H3K27ac signal is better capable of distinguishing different PDC cell-states, suggesting that active enhancer and promoter landscapes are more unique to each individual cell state when compared to just promoter activity alone. Furthermore, our results indicate that gain of H3K27ac at promoters and/or poised enhancers may serve as critical factors driving tumour progression. We validated that H3K27ac/H3K4me3 activity at the KRT5 and KRT8/18 loci served as an accurate indicator of epithelial identity, and that snCUT&RUN can be used to comprehensively analyze a mixture of samples to infer epithelial changes and the accompanying TF motif changes during tumour progression. Importantly, we showed that these changes are possibly associated with a focal copy number amplification of YAP1, which in turn could lead to altered TEAD activity and subsequent epigenetic rewiring. Indeed, an increasing body of evidence show that genetic and epigenetic factors are closely associated with driving tumour heterogeneity and progression. An example of this interplay between the genome and the epigenome was highlighted in recent studies on extrachromosomal circular DNA (ecDNA) amplifications that not only amplify oncogenes, but also functional elements such as enhancers, thereby increasing chromatin accessibility and downstream expression of key oncogenes (Wu et al. 2019; Morton et al. 2019). We further showed that gene expression is affected by both gene copy number as well as gene chromatin state, where the highest gene expression was observed in genes that have higher copy number *and* have active chromatin marks. These results emphasize the complexities of factors that promote tumour progression and highlight the importance of considering both genetic and epigenetic factors that drive cell-state plasticity.

However, we noted that some epigenetic changes were more subtle and did not involve wide-scale epigenetic rewiring driven by alterations in the genome. Comparing both HN120Pri and HN120Met as well as HN137Pri and HN137PCR, we did not observe clear lineage infidelity from both H3K4me3 and H3K27ac profiles. This led to the hypothesis that in some tumours, microenvironmental pressures such as metabolic dysregulation or challenges from the immune system may lead to primary tumour subpopulations to activate certain transcription factor programmes to adapt to these selective pressures. This adaptation process inherently leads to heterogeneous chromatin states in the primary tumour, which may confer selective fitness to the subpopulations with an intermediate malignant chromatin state. Indeed, a previous study described a subpopulation of malignant HNSCC cells being in a state of pseudo-epithelial to mesenchymal transition (pseudo-EMT), which correlated with cancer progression (Puram et al. 2017). We used the module score analysis and showed that H3K27ac profiles could be used to find such subpopulations that may have a higher propensity to progress to malignant cell states.

We note that the samples we have used for this study are long-term cell cultures which may have acquired a degree of homogeneity through the course of several passages. However, some evidence suggests that heterogeneity is maintained even in long-term cultured cell lines. For example, our cell line models exhibit chromosomal instability (CIN). CIN is known to lead to heterogeneity in gene expression, while at the same time being maintained in long term cell cultures. In a study by Minussi et al. (Minussi et al. 2021), the authors performed single-cell clonal outgrowth of the CIN-affected MDA-MB-231 breast cancer cell line and found that karyotype diversity was regained after only 19 passages, suggesting that *in vitro* cancer cell cultures re-diversify their genomes and maintain genetic heterogeneity throughout passaging. Epigenomic heterogeneity was also shown to be maintained in cell lines. Litzenburger et al. observed through scATAC-seq experiments that the leukemic cell line K562 exhibited epigenomic heterogeneity which has functional consequences such as drug response (Litzenburger et al. 2017). Hence, it is likely that our cell line models exhibit sufficient intratumour heterogeneity to make single-cell assessments useful.

Altogether, based on the observations from snCUT&RUN analysis on isogenic primary and progressed cell states in HNSCC, we propose the following working model for epigenetic control of HNSCC progression (Fig. 4F). First, in the presence of a strong genetic driver, e.g., a copy number amplification, epigenetic reprogramming can occur, which may result in the selection of cells with favorable gene expression signatures to progress. Second, in the absence of strong genetic drivers, microenvironmental pressures may drive a primary tumour adaptation response, leading to intratumour epigenetic heterogeneity. ITeH results in the emergence of transitionary subpopulations that are more prone to progress into a more malignant or resistant cell-state. Future characterization of such sub-populations can result in the identification of prognostic biomarkers and therapeutic targets that can be used for better patient stratification and the development of novel intervention strategies to block progression to malignant states.

At present, snCUT&RUN is positioned as a medium-throughput assay. Slightly more than 2000 FACS sorted single cells were profiled in this work. Although it may not achieve the throughput of scCUT&TAG assays, its high sensitivity is a major advantage for uncovering subtle cell-to-cell epigenetic heterogeneity, such as those found within a tumour. We envisage that future improvements such as automation of single cell library preparation with an automated liquid handler would further enable snCUT&RUN assay to be adopted by the wider scientific community.

## Methods

### Cell materials and tissue culture

HN120Pri, HN120Met, HN137Pri and HN137Met patient derived cells (PDCs) used for this study were retrieved from a biobank previously generated using described methods (Chia et al. 2017). Likewise, HN120PCR and HN137PCR cells were generated as part of an earlier study (Sharma et al. 2018). The identities of the primary cell lines and their derivative were confirmed using short-tandem repeat (STR) profiling (Supp. Table 1). Normal tissue samples for these patients were not available for experiments. Cells were cultured with RPMI medium supplied by 10% Fetal Bovine Serum (FBS) and 1% penicillin/streptomycin. Medium was replaced every 2-3 days. Cells were cultured in 37°C and 5% CO_2_ and passaged when cultures reached ∼90% confluency. Cells were tested for mycoplasma and were only used for experiments after being confirmed to be mycoplasma negative.

### Immunofluorescence

Cells were cultured in 96-well plates with 10,000 – 20,000 seeding density for 48 hours. Fixation was done in acetomethanol 1:1 ratio for 10 minutes at −20°C degrees. After washing 3X with 1X PBS, blocking was done with 2% BSA/0.1% Triton X-100/PBS for 1hr+. The following primary antibodies were used: ab17130 (KRT5, 1:100 dilution) and ab32118 (KRT18, 1:100 dilution), and cells were incubated at 4°C overnight. Alexa Fluor488/594 and Hoechst33342 were used during secondary antibody incubation for 30 minutes at 37°C. Cells were imaged with the Nikon EclipseTi inverted microscope using widefield setting and 20X magnification.

### Single nuclei CUT&RUN

Cells cultured to ∼80% confluence in T75 flasks were harvested and washed 2X in Sucrose Buffer (5% sucrose, 1% BSA, 20mM HEPES pH7.5, 150mM sodium chloride 0.5mM spermidine, 1X cOmplete™, Mini, EDTA-free Protease Inhibitor Cocktail (Roche, Cat# 04693159001) at 200 x g for 3 min. For each antibody tested, 1.5 x 10^6^ cells were dispensed into Protein LoBind® Tubes (Eppendorf, Cat# 0030108442), pelleted and suspended in 1X ice cold lysis buffer (Sucrose Buffer containing 0.005% NP-40, 0.005% digitonin, 2mM EDTA and 10mM sodium butyrate). The amount of cell lysis was detected using the trypan blue exclusion assay, and remaining intact cells were lysed by titrating-in 2X cold lysis buffer (Sucrose Buffer containing 0.01% NP-40, 0.01% digitonin, 2mM EDTA and 10mM sodium butyrate). Nuclei were pelleted then suspended in Sucrose Buffer containing 1:100 anti-Histone H3 (tri methyl K4) antibody (abcam, Cat# ab213224) or anti-acetyl-Histone H3 (Lys27) (Merck, Cat# MABE647) and incubated ≥ 1.5h at 4°C with intermittent agitation. Nuclei were washed 2X with Sucrose Buffer then suspended in 700 ng/ml pA-MN (a kind gift from Steven Henikoff) in Sucrose Buffer and incubated for 60 min at 4°C. Pelleted nuclei were suspended in Nuclear Stain Buffer (Sucrose Buffer containing 1:4,000 Alexa Fluor® 647 anti-Nuclear Pore Complex Proteins Antibody (BioLegend, Cat# 682204) plus 4µM ethidium homodimer (Invitrogen, Cat# E1169) and incubated at room temperature (RT) for 10 min. After 2X washes with Sucrose Buffer the nuclei were suspended in Low Salt Buffer (20 mM HEPES pH 7.5, 0.5 mM spermidine). FACS was performed with an MoFlo Astrios Cell Sorter (Beckman Coulter) operated using Summit software. Single nuclei were gated first using forward and side scatter pulse area parameters (FSC-A and SSC-A), aggregates excluded using pulse width (FSC-W and SSC-W), then isolated nuclei were gated based on AF647 and ethidium homodimer fluorescence. Nuclei were sorted directly into 3 µl of calcium buffer (10 mM calcium chloride, 3.5 mM HEPES pH 7.5) in PCR strip tubes for DNA digestion. As a quality control measure, nuclei were also sorted into buffer on flat-well optical plates (4titude, Cat# 4ti-0970/RA) to check for wells with more than one isolated nuclei via fluorescence microscopy. For each array of 96 PCR tubes, positive and negative control tubes were included which comprised of 1,000 dispensed nuclei or buffer-only wells, respectively. To stop digestion, 1 µl of 4X STOP Buffer (600 mM sodium chloride, 80 mM EGTA and 0.05% digitonin) was added to the sides of the strip tubes, the tubes pulse centrifuged, mixed by touching the sides of the tubes against a rotating vortex, then pulse centrifuged a second time. Tubes were then incubated at 37°C for 30 min.

Library prep on single nuclei was performed as follows: 0.8 µl 1X End Repair and A-Tailing Buffer/Enzyme Mix (KAPA Hyper Prep Kit, Cat# 07962363001) was added to the sides of the tubes, mixed as before, then incubating at 12°C for 15 min, 37°C for 15 min, 58°C for 90 min, then 8°C on hold. Adapter ligation was performed by adding 0.5 µl 300 nM Unique Dual Indexed adapter (NEXTFLEX, Cat# NOVA-514150 or NOVA-514151) and 3.6 µl 1X Ligation Buffer/Enzyme mix (KAPA Hyper Prep Kit, Cat# 07962363001) to the sides of the tubes, mixed as before, then incubating for 16 hours at 4°C.

To remove excess adapters, the volumes were adjusted to 20 µl with 10 mM Tris pH 8.0, 20 µl 2X SPRI beads (MagBio Genomics, Cat# AC-60050) [2X SPRI beads: 1 volume SPRI beads magnetically separated, ½ volume PEG/NaCl solution removed (and stored), then beads re-suspended. This increases the surface area of beads for DNA to adsorb to.] added to inverted fresh strip tube caps, the caps were then affixed, the tubes mixed as before then incubated at RT for 2 hours. SPRI beads were separated by tethering the strip tubes to N50 grade neodymium magnets affixed to a ferrous metal rig for stability. The supernatant was removed, the beads washed twice with 80% ethanol then allowed to dry for 3 min at RT. The tubes were removed from the magnets and beads suspended in 20 µl 10 mM Tris pH 8.0. 22 µl PEG/NaCl was added and mixed into the beads with pipetting, then incubated for 2 hours at RT (or tubes left overnight at 4°C, brought back to RT, then incubated for 2 hours at RT). Beads were washed twice with 80% ethanol, air dried for 3 min then suspended in 7.5 µl 10 mM Tris pH 8.0 and DNA allowed to elute for ≥ 10 min at RT. Hereafter the positive control tubes were treated separately and not pooled with single nuclei libraries and no template controls. The beads were magnetically separated as above and pooled eluate added to PCR strip tubes containing 2X KAPA HiFi HotStart ReadyMix (Roche, Cat# 07958935001) and 2 mM each of P5 (AATGATACGGCGACCACCGAGATCTACA*C) and P7 (CAAGCAGAAGACGGCATACGAGA*T) primers with phosphorothioate bond as indicated with asterisk. PCR was performed with a thermocycler using a heated lid under the following cycling conditions: 98°C for 45s; 19 cycles of 98°C for 15 s, 60°C for 10 s; 72°C 1 min, 8°C on hold.

Pooled library DNA was concentrated as follows: amplified DNA was pooled together, 400 µl aliquots dispensed into 1.5 ml tubes, 400 µl Phenol:Chloroform:Isoamyl Alcohol 25:24:1 added and the samples vortexed. The mixture was then transferred to MaXtract High Density tubes (QIAGEN, Cat# 129056) and centrifuged at RT for 5 min at 16,000 x g. 400 µl chloroform was then added to the MaXtract tubes, mixed by inverting several times, then centrifuged at RT for 5 min at 16,000 x g. The aqueous phase from each MaXtract tube (∼400 µl) was then transferred to 1.5 ml tubes containing 2 µl of 2 µg/µl glycogen (Roche, Cat# 10901393001), 1 ml ethanol was added, the tubes vortexed, then incubated at −20°C overnight. DNA was pelleted by centrifuging at 20,000 x g, 10 min at 4°C, washed with 1 ml 100% ethanol, then air dried for at least 15 min at room temperature. Pellets were suspended and combined in a total volume of 100µl 10 mM Tris pH 8.0. Adapter dimers and excess primers were removed by adding 90 µl SPRI beads and incubating for 15 min at RT, magnetically separating the beads, washing twice with 80% ethanol and eluting with 50µl 10 mM Tris pH 8.0. DNA was allowed to rebind to the beads by adding 45 µl PEG/NaCl, the beads were incubated for 15min at RT, magnetically separated, washed twice with 80% ethanol, then eluted in 50 µl 10 mM Tris pH 8.0. A third SPRI bead purification with a ratio of 1:1 was found to be necessary to remove residual adapter dimers prior to sequencing (i.e. 50 µl PEG/NaCl added to the DNA/bead mixture, and two ethanol washes performed). Finally, the DNA was eluted in 20 µl 10 mM Tris pH 8.0. All libraries were sequenced using paired-end sequencing on an Illumina MiSeq™, with samples processed using MiSeq Reagent Kit V3 (150 cycles) (Illumina, Cat#MS-102-3001).

### Bulk cell CUT&RUN

Bulk cell CUT&RUN was performed in the same manner as snCUT&RUN with the following alterations. After incubating with pA-MN, cells were washed twice in BSB followed by one wash in Low Salt buffer. After pelleting, most of the supernatant was removed leaving a small volume and the pellet suspended to a slurry by gentle agitation. 37 µl calcium buffer was added to activate pA-MN and the samples incubated in a pre-chilled heat-block in ice-water for 30 min. The reaction was stopped by adding 12.5 µl 4X STOP Buffer and nucleosomes allowed to diffuse for 30 min at 37°C. 50 µl was removed to a PCR tube containing 10 µl 1X End Repair and A-Tailing Buffer/Enzyme Mix (KAPA Hyper Prep Kit, Cat# 07962363001), then incubated at 12°C for 15 min, 37°C for 15 min, 58°C for 90 min, followed by 8°C on hold. 5 µl 15 mM TruSeq DNA Single Indexes (SKU 20015960) and 45µl 1X KAPA Hyper Prep Enzyme/Buffer mix were added to each sample and incubated for 16 hours at 4°C. 2 µl 10% SDS plus 2 µl proteinase K (Thermo Fisher Scientific, Cat# EO0492) were added and incubated for 60 min 37°C. Excess adapters and adapter dimers were removed via two successive washes with SPRI beads. 110 µl SPRI beads (MagBio Genomics, Cat# AC-60050) were added to each sample and incubated for 15 min at room temperature. Beads were magnetically separated and washed twice with 80% ethanol. After air drying for 5 min, DNA was eluted from the beads by suspended in 50 µl 10 mM Tris pH 8.0. DNA was re-bound to the beads by adding 60 µl 20% PEG 8000, 2.5 M NaCl and incubating for 15 min. Beads were separated and washed as before, then suspended in 20 µl 10 mM Tris pH 8.0. To each purified library, 25 µl 2X KAPA HiFi HotStart ReadyMix added plus 5 µl TruSeq single index PCR Primer Cocktail. PCR was performed using the following cycling conditions: 98°C for 45s; 12 cycles of 98°C for 15 s, 60°C for 10 s; 72°C 1 min, 8°C on hold. Bulk cell libraries were purified using two successive rounds of SPRI bead purification as above, using 1:1.1 then 1:1.2 ratios of sample to beads, and eluting with 20 µl 10 mM Tris pH 8.0.

### snCUT&RUN data preprocessing and QC measurements

Raw .fastqs of single-cells were mapped to the human hg38 reference genome with bowtie2 (Langmead and Salzberg 2012, 2) v2.3.5.1 and the following settings: --end-to-end --very-sensitive --no-mixed --no-discordant -q --phred33 -I 10 -X 700. The MarkDuplicates tool from GATK v4.1.4.1 (McKenna et al. 2010) was used to mark and remove duplicate reads while samtools v1.10 (Li et al. 2009) was used to index the bam files. Bedtools v2.27.1 (Quinlan and Hall 2010) was used to convert .bam files to .beds and .bedgraphs. The unique number of UMRs was calculated by first converting the deduplicated single cell .bam file to a .bed file, which in turn was used to create a TagAlign file. The number of paired-end reads in the TagAlign file was then counted as the number of UMRs. To calculate the fraction reads in peaks, we first merged the single-cell data into pseudobulk, and then normalized based on the sample with the lowest read number. We then called peaks using MACS2 with the following settings: BAMPE --nomodel -B -p 5e-2 --min-length 500 --max-gap 400 –SPMR --call-summits. The fraction of reads that overlap the called peaks was then divided by the total number of reads to calculate the FRiP. Equally, we calculated the fraction of peaks in blacklisted genomic regions (Amemiya et al. 2019).

### Signac/Seurat analysis

To create Signac fragment files, individual single-cell .bam files were first processed into .bed files with the following columns: chr, start, end, cell barcode, and a fifth column with the number 1, representing that each row combination only occurred once. Individual bed files were concatenated and sorted with bedtools and indexed with tabix. Data from H3K4me3 and H3K27ac were processed separately. The fragment file was then converted to a sparse bin matrix with Signac’s GenomeBinMatrix() function in R4.0, specifying binsize of 10kb. Subsequently, the bin matrix was used to create a chromatin assay in Signac. Cells were excluded from analysis with the following criteria: UMRs < 1,000; UMRs > 100,000; FRiP < 0.25; fraction reads in blacklist > 0. 01; nucleosome signal > 5 and TSS enrichment score < 0.5. Signac standard analysis was performed downstream. Briefly, the matrix was normalized with term frequency inverse document frequency (TF-IDF), and singular value decomposition (SVD) was applied. UMAP embedding was done with the following settings: umap.method = “uwot”, n.neighbors = 10, metric = “manhattan”, n.epochs = 500, min.dist = 0.1, spread = 1, set.op.mix.ratio = 1, reduction = ‘lsi’, dims = 2:30. MACS2 peaks were called with the CallPeaks() function, specifying the broad and combine.peaks options set to TRUE, BAMPE --nomodel -B -p 5e-2 --min-length 500 --max-gap 400 --SPMR --call-summits. Peak calls were visualized with the CoveragePlot() function. To perform motif analysis, first a PFMatrix object was retrieved from the JASPAR2020 package (Fornes et al. 2020). A peak matrix was then created using the fragment files and peak calls. Subsequently the AddMotifs() function was used to construct a motif object containing motif information. ChromVAR deviation Z-scores were calculated with the RunChromVAR function. Reactome pathway analysis of peaks was done with the rGREAT package v1.24.0 (McLean et al. 2010; Gu 2022). For this analysis, only peaks with differential p-value of <0.005 were included.

### Bulk RNA-seq

Total RNA was extracted with the Qiagen RNAeasy Plus Mini Kit (Cat. No. 74136) and quantified with NanoDrop. Triplicates for each cell line (using different passage number) were used. RNA-seq library preparation and directional mRNA-sequencing was carried out by NovogeneAIT. And adapted analytical workflow from (Love et al. 2019) was used for the analysis. Briefly, transcripts were quantified using Salmon (Patro et al. 2017) with hg38 genome and GENCODE version 38 as reference and default parameters. Quantified transcripts were imported to R with tximeta (Love et al. 2020) and differential analysis was performed with DESeq2 (Love et al. 2014), filtering out genes with less than 10 supporting reads. Z-scores were calculated with the scale() function after variance normalization with variance stabilizing transformation (VST) to account for count variations caused by highly expressed or lowly expressed genes. The resulting output is a matrix where rows represent genes, columns represent samples, and values represent Z-score. The mean Z-score of the three replicates was calculated as the representative Z-score for each individual PDC.

### Whole exome sequencing

WES data for HN120Pri, HN120Met, HN137Pri and HN137Met were done through Macrogen as part of an earlier study (in press), while WES for HN120PCR and HN137PCR was done through NovogeneAIT. In this case, the Agilent SureSelect V6 58 Mb kit was used for library preparation and samples were sequenced on an Illumina NovaSeq PE150 platform at 12Gb data/100X exome coverage. Raw WES data were processed through GATK best practices pipeline, and copy number segmentation and calling were done with CNVKit after normalizing read number to the lowest sample (HN137Met) (Talevich et al. 2016).

### Correlating gene copy number, gene chromatin state, and gene expression

Single cell .bams were first aggregated into pseudobulk with samtools merge and processed with sorting and indexing. Next, to account for differences in signal arising due to differences in read number, we normalized the .bam files by downsampling reads to match the read number of the sample with the smallest read number. Normalized bams of pseudobulk aggregate H3K4me3 and H3K27ac, as well as bulk H3K27me3 data were used as input for ChromHMM v1.23 (Ernst and Kellis 2012), using hg38. ChromHMM is a multivariate Hidden Markov Model-based technique that can model the presence of multiple chromatin marks in the same region of the genome. It first uses the Baum-Welch algorithm to find ‘hidden chromatin states’, each state reflecting presence or absence of single or multiple histone marks. It then uses the forward-backward algorithm to calculate the posterior probability of a certain genomic region being in a particular chromatin state. Five states were deemed to be the optimal number of states after repeated analysis with multiple number of states. Output segment files were binned to 200bp windows with bedtools makewindows to allow for comparable analysis across multiple cell lines. To correlate ChromHMM state transitions and gene expression, ChromHMM 200bp genomic bin outputs were filtered to include only promoter regions (region of 2000bp upstream of TSS to 3000bp downstream of TSS) and correlated to the nearest gene with bedtools closest. The resulting output is a bed like file with the columns: chromosome, bin start, bin end, chromatin state, and gene. Gene symbols were used as identifier to connect gene chromatin state, gene copy number, gene expression. The Z-score matrix and CNVKit .call.cns output was used for the gene expression and gene CN data respectively. Genes with missing data (e.g. no gene expression or CNV data) were removed from analysis. To assign chromatin state from the 200bp ChromHMM output, we considered a gene to have an activating mark if more than 2 bins (400bp) have H3K4me3 signal, H3K27ac signal, or both. Genes were categorized as having activity of both marks if the modal chromatin state across all bins are H3K4me3+/H3K27ac+ or when there are similar proportion of H3K4me3+/H3K27ac- and H3K4me3-/H3K27ac+ bins (e.g. 40% H3K4me3+/H3K27ac- and 60% H3K4me3-/H3K27ac+ or vice versa). A gene is categorized as having single mark if more than 60% of the bins were annotated as such by ChromHMM. For chromatin state transition analysis, bins that were unmodified in the primary state and which remained unmodified during progression were excluded from analysis. Alluvial plots were plotted with the R package ggalluvial (Brunson 2020). Barplots and boxplots were visualized with the R package ggplot2.

### HN120preMet and HN137prePCR analysis

Differential marker peaks for each individual PDC were found through the FindAllMarkers() function of Seurat. The top-50 peaks were included as features for the “module” unique for each PDC. Subsequently, module scores were back calculated for each single cell with the AddChromatinModule() function. To plot the ternary plots, first individual module profiles of HN120 and HN137 PDCs were separated. Next, normalized module score profiles of each individual cell were added up to 1 with the following formula: 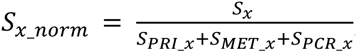, whereby S_x_ is the raw PRI or MET or PCR module score of that cell, and S_(x_norm)_ is the normalized PRI or MET or PCR score for that cell. If there were negative values, the absolute value of the lowest negative value was added to the PRI/MET/PCR scores. The package ggtern (Hamilton and Ferry 2018) was then used to visualize the normalized module scores. For HN120Pri, cells were categorized as preMet if cells have a PRI/MET/PCR normalized module score ratio of >0.4 Pri and >0.4 Met. For HN137Pri, cells were relabeled as HN137prePCR if cells have a PRI/MET/PCR normalized module score ratio of >0.4 Pri / <0.4 PCR. The ggplot package and the geom_smooth() function was used to calculate the linear regression between HN120Pri and HN120Met module scores with UMR and FRiP.

### Statistical analysis

All statistical analysis was done with R4.0. The two-sided Wilcoxon test from the stat_compare_means function of the ggpubr package was used to calculate statistical significance of either module scores, UMRs, or FRiP between different PDCCs. A linear regression model was used to analyze the statistical relationship between HN120Met module score and FRiP and UMR. P-values as indicated: *, p <= 0.05; **, p <=0.01; ***, p <= 0.001; ****, p <= 0.0001.

## Data access

The datasets supporting the conclusions of this article are available in the NCBI GEO repository, under the accession number <u>GSE212250</u> (https://www.ncbi.nlm.nih.gov/geo/query/acc.cgi?acc=GSE212250). The code used in this study are available on Github: https://github.com/dmuliaditan/sncutnrun. Figures 1A and 4F were created with Biorender (www.biorender.com) under publication license HI248I94WF and GS26ET99NE respectively.

## Acknowledgments

We thank the members of the DasGupta and Cheow labs for helpful discussion. We also thank Xiaoning Wang from the NUS Medicine Flow Cytometry Laboratory for assistance and advice on cell sorting and analysis. This work is supported by a grant from the National Medical Research Council (Singapore) (MOH-000219).

## Author’s contributions

Conceptualization: HJW, RD, LFC. Methodology: HJW, DM. Supervision: RD, LFC. Writing: HJW, RD, RD, LFC.

## Author Information

Howard J. Womersley and Daniel Muliaditan are co-first authors of this study.

